# Distribution and genomic variation of thermophilic cyanobacteria in diverse microbial mats at the upper temperature limits of photosynthesis

**DOI:** 10.1101/2022.03.25.485844

**Authors:** Eric D. Kees, Senthil K. Murugapiran, Annastacia C. Bennett, Trinity L. Hamilton

## Abstract

Thermophilic cyanobacteria have been extensively studied in Yellowstone National Park (YNP) hot springs, particularly from decades of work on the thick laminated mats of Octopus and Mushroom Springs. However, focused studies of cyanobacteria outside of these two hot springs have been lacking, especially regarding how physical and chemical parameters along with community morphology influence the genomic makeup of these organisms. Here, we used a metagenomic approach to examine cyanobacteria existing at the upper temperature limits of photosynthesis. We examined 15 alkaline hot spring samples across six geographic areas of YNP, all with varying physical and chemical parameters, and community morphology. We recovered 22 metagenome-assembled genomes (MAGs) belonging to thermophilic cyanobacteria, notably an uncultured *Synechococcus*-like taxon recovered from the upper temperature limit of photosynthesis, 73°C, in addition to thermophilic *Gloeomargarita*. Furthermore, we found that three distinct groups of *Synechococcus*-like MAGs recovered from different temperature ranges vary in their genomic makeup. MAGs from the uncultured very high temperature (up to 73°C) *Synechococcus*-like taxon lack key nitrogen metabolism genes and have genes implicated in cellular stress responses that diverge from other *Synechococcus*-like MAGs. Across all parameters measured, temperature was the primary determinant of taxonomic makeup of recovered cyanobacterial MAGs. However, Fe, community morphology, and biogeography played an additional role in the distribution and abundance of upper temperature limit-adapted *Synechococcus*-like MAGs.These findings expand our understanding of cyanobacterial diversity in YNP and provide a basis for interrogation of understudied thermophilic cyanobacteria.

**Importance:** Oxygenic photosynthesis arose early in microbial evolution – approx. 2.5-3.5 billion years ago – and entirely reshaped the biological makeup of Earth. However, despite the span of time in which photosynthesis has been refined, it is strictly limited to temperatures below 73°C, a barrier that many other biological processes have been able to overcome. Furthermore, photosynthesis at temperatures above 56°C is limited to circumneutral and alkaline pH. Hot springs in Yellowstone National Park (YNP), which have a large diversity in temperatures, pH and geochemistry provide a natural laboratory to study thermophilic microbial mats, and the cyanobacteria within. While cyanobacteria in YNP microbial mats have been studied for decades, a vast majority of work has focused on two springs within the same geyser basin, both containing similar community morphologies. Thus, the drivers of cyanobacterial adaptations to the upper limits of photosynthesis across a variety of environmental parameters have been understudied. Our findings provide new insights into the influence of these parameters on both taxonomic diversity and genomic content of cyanobacteria across a range of hot spring samples.

## Introduction

Oxygenic photosynthesis in Cyanobacteria is among the most impactful microbial innovations in Earth’s history and accounts for the largest source of O_2_ in the atmosphere. While photosynthesis has had approximately 2.5-3.5 billion years of evolution (Cardona et al., 2019; Fischer et al., 2016; Martin et al., 2018), it is constrained to temperatures < 73°C in alkaline environments and < 56°C in acidic environments (Boyd et al., 2010; Brock, 1973; Brock & Brock, 1968; Hamilton et al., 2012; Toplin et al., 2008).

Characterizations of high temperature cyanobacteria have been most extensively done in Hunter’s Hot Springs, OR, USA and in thick laminated microbial mats of alkaline Mushroom Spring, Octopus Spring, and the mat and streamer communities in White Creek (reviewed in Ward et al., 2012) in Yellowstone National Park (YNP), WY, USA hot springs. A limited number of studies have focused on phototrophic hot spring communities elsewhere in YNP (Becraft et al., 2020; Bennett et al., 2020; Fecteau et al., 2022; Hamilton et al., 2019; Inskeep et al., 2013). The most prevalent cyanobacteria found at or near the 73°C limit in these springs are those in a *Synechococcus*-like clade (Allewalt et al., 2006; Ferris & Ward, 1997; Ward et al., 2006). Since these thermophilic *Synechococcus*-like cyanobacteria form a basal clade, distantly related to all other known *Synechococccus* species, they were recently classified under the proposed genus name *Leptococccus* (Salazar et al., 2020; Walter et al., 2017), or more recently *Thermostichus* (Komárek et al., 2020). For the purposes of clarity and contextualization around existing literature, we refer to these cyanobacteria in the present study as belonging to the *Synechococcus-*like A/B lineage (Allewalt et al., 2006; Becraft et al., 2011, 2020; Ferris & Ward, 1997; Papke et al., 2003; Ramsing et al., 2000; Ward et al., 1990).

Molecular probing of *Synechococcus*-like cyanobacteria in Octopus and Mushroom Springs revealed several “ecotypes”, or sequence variants occupying different niche spaces defined by environmental variables such as temperature and light (Becraft et al., 2011; Bhaya et al., 2007; Ferris & Ward, 1997; Gomez-Garcia et al., 2011; Ramsing et al., 2000; Ward et al., 1990). These ecotypes were organized into major categories designated “A” and “B”. “A” ecotypes (including the isolated and sequenced strain JA-3-3Ab, referred to here and elsewhere as OS-A), tend to occupy the highest temperatures up to 73°C, while “B” ecotypes (including JA-2-3B’a(2-13) or OS-B’) occupy a slightly lower range up to 65°C (Allewalt et al., 2006; Bhaya et al., 2007; Ferris & Ward, 1997). A and B ecotypes have been more finely delineated along temperature gradients into A, A’, A’’, B’ and so on, generally with each “prime” designation indicating a temperature range trending upward (Ferris & Ward, 1997). Similar niche differentiation along temperature gradients was described for *Synechococcus*-like cyanobacteria in Hunter’s Hot Springs and at present, the only cultured A’-like cyanobacteria were isolated from this location at 70 °C (Miller et al., 2013; Miller & Carvey, 2019; Miller & Castenholz, 2000; Pedersen & Miller, 2017). Biogeography plays an additional role in genetic divergence of ecotypes within A/B-lineage clades (Becraft et al., 2020). In the present study, we were primarily interested in the genetic factors that drive ecological diversification among A/B-lineage populations, particularly in YNP hot springs with differing pH, temperature, and geochemistry.

While the underlying genetic factors driving adaptation to different temperatures among *Synechococcus*-like A/B-lineage organisms remain largely unresolved, some specific adaptations to the highest temperatures have been identified. For example, more thermostable variants of RuBisCO have been found to extend the thermal niche in an A/B-lineage clade from Hunter’s Hot Springs (Miller et al., 2013). Comparisons between OS-B’ (53-60°C) and OS-A (58-70 °C) isolates from Octopus Spring also revealed different strategies for phosphorus assimilation: in contrast to OS-B’, OS-A lacks a C-P lyase operon and is unable to grow heterotrophically in the dark from phosphonates (Gomez-Garcia et al., 2011). Since focused studies of high temperature *Synechococcus*-like A/B-lineage ecotypes have been largely limited to the thick laminated mats of Octopus, Mushroom, and Hunter’s Hot Springs, it is unknown whether they are driven by the same adaptations in other springs. Specifically, the degree to which pH influences temperature adaptation and taxonomic makeup of A/B-lineage cyanobacteria is understudied.

Here we leveraged metagenomic sequencing of samples across a range of pH, temperature, geochemistry, and community morphology to further define the taxonomic distribution and genomic content of the A/B-lineage and other thermophilic cyanobacteria within YNP hot springs. We focused specifically on communities existing in a range of temperatures at circumneutral to alkaline pH, and in communities with morphologies ranging from thin and thick mats to filaments. We asked whether the niches defined for the *Synechococcus*-like A/B-lineage in Mushroom and Octopus springs are applicable to other Yellowstone hot springs, and whether the distribution and relative abundances of phototrophic taxa are constrained by a combination of temperature and pH. Using a pangenome approach, we detail genomic comparisons of 15 *Synechococcus*-like A/B-lineage metagenome-assembled genomes (MAGs) from three distinct clades.

## Results and discussion

### Sample site descriptions and recovery of metagenome assembled genomes

We asked whether the temperature constraints placed on laminated, cyanobacterial mats in alkaline Mushroom and Octopus springs were similar in other Yellowstone hot springs with varying geography, community morphology, and pH. Additionally, we examined the extent to which pH constrains the upper temperature limit and taxonomic composition of thermophilic cyanobacteria. We collected 16 samples from 11 hot springs within 3 geyser basins in 2017 and 2018 (Fig 1, Table 1), targeting the transition between pigmented photosynthetic communities and chemotrophic zones, termed the photosynthetic “fringe”. YNP regions sampled include the Middle Geyser Basin (Rabbit Creek Area), Lower Geyser Basin (Boulder Geyser; Imperial Geyser Basin; Sentinel Meadows; White Creek Area), and Gibbon Geyser Basin (Geyser Creek Area). Generally, sample temperatures ranged from 44.2°C to 73.0°C, and pH ranged from 7.3 to 9.4. Sulfide concentrations ranged from 0.22 µM to 4.93 µM (with one exception at 18.99 µM), chloride from 7.7 µM to 12.0 µM, and iron from 93.8 nM to 765.3 nM, In samples taken from 2018 nitrate concentrations were detectable up to 14.3 µM, while ammonium concentrations were detected up to 6 µM. No fixed nitrogen data was available for 2017 samples.

**Figure 1.**
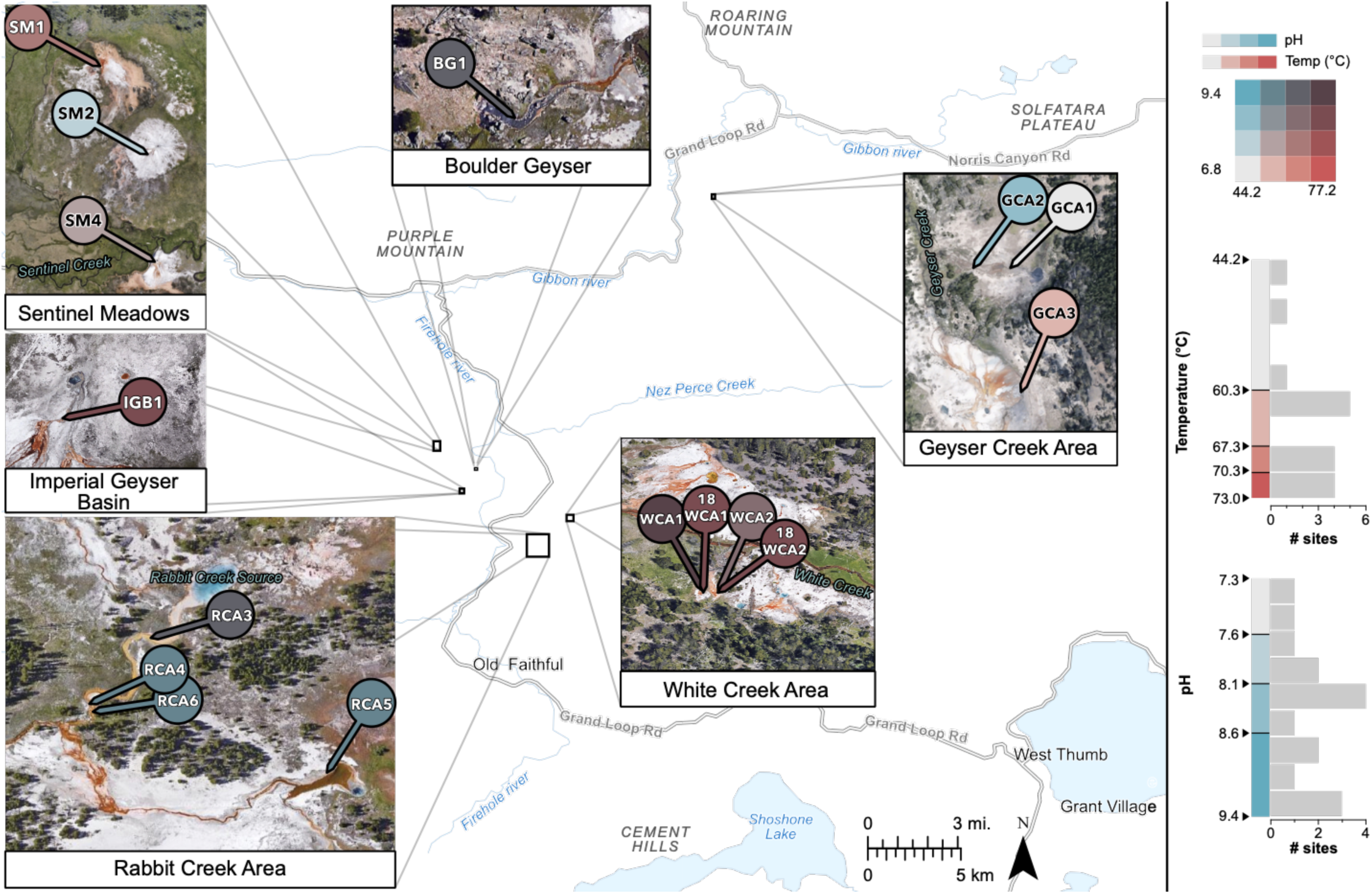
Sample Locations. Samples were collected in Yellowstone National Park at locations with varying temperatures, pH, and community morphology. Samples are shown sorted into 4 temperature (red) and 4 pH (blue) color coordinates. Sorting was done according to geometric intervals to account for skewed data and better differentiate temperatures near the upper limits.

**Table 1.** Metagenome Sample descriptions. RCA=Rabbit Creek Area ; SM = Sentinel Meadows ; BG = Boulder Geyser ; WCA = White Creek Area ; GCA = Geyser Creek Area ; IGB = Imperial Geyser Basin.

| Sample Name | Sample ID | IMG Genome ID | BioSample Acc. | Size (bp) | Gene count | pH | T (°C) | Community Morphology |
| --- | --- | --- | --- | --- | --- | --- | --- | --- |
| BG1 | 170629B | 3300028893 | SAMN26200800 | 196918189 | 436827 | 8.7 | 68.5 | Red mat and filaments |
| GCA1 | 170630B | 3300038630 | SAMN26200803 | 222603420 | 441247 | 7.6 | 59.9 | Orange/yellow |
| GCA2 | 170630C | 3300040982 | SAMN26200804 | 291888643 | 632804 | 8.3 | 44.2 | Pink-white-grey mat |
| GCA3 | 170630D | 3300028820 | SAMN26200805 | 213192026 | 395775 | 7.3 | 62.7 | Red/grey mat |
| IGB1 | 180603O | 3300036711 | SAMN26200807 | 117128738 | 264866 | 8.2 | 66.6 | Orange/yellow mat |
| RCA3 | 170626A | 3300028606 | SAMN26200794 | 114093415 | 252304 | 9.1 | 68.4 | Thin Yellow/orange mat |
| RCA4 | 170626B | 3300028609 | SAMN26200795 | 195353357 | 412629 | 9.2 | 62.3 | Orange/yellow mat |
| RCA5 | 170626C | 3300028611 | SAMN26200796 | 108943318 | 208402 | 9.4 | 62.4 | Orange/yellow mat |
| RCA6 | 170626D | 3300028818 | SAMN26200797 | 182682112 | 386452 | 9.3 | 62.3 | Orange/yellow mat |
| SM1 | 170627A | 3300028613 | SAMN26200798 | 316999032 | 710899 | 7.9 | 68.4 | Thin yellow, orange mat |
| SM2 | 170627E | 3300028839 | SAMN26200799 | 475303863 | 1088432 | 8.1 | 51.4 | Orange/yellow mat |
| SM4 | 180602L | 3300038980 | SAMN26200806 | 76631469 | 142361 | 8.0 | 62.3 | Orange/yellow mat |
| WCA1 | 170629F | 3300028816 | SAMN26200801 | 104661654 | 234717 | 8.8 | 71.0 | Green filaments |
| 18WCA1 | 180603R | 3300038981 | SAMN26200808 | 101090071 | 215532 | 8.4 | 73.0 | Green filaments |
| WCA2 | 170629H | 3300028617 | SAMN26200802 | 131707792 | 292784 | 8.6 | 69.4 | Green filaments and mat |
| 18WCA2 | 180603S | 3300038484 | SAMN26200809 | 142901704 | 282903 | 8.2 | 71.9 | Green filaments |

In the Middle Geyser Basin, samples taken from the Rabbit Creek Area (RCA) consisted of yellow/orange mats. Samples RCA3, RCA4, and RCA6 were collected from Rabbit Creek a fast-moving outflow in a length along the channel spanning ∼ 70-150 m from the source spring and between 68.4 and 62.3 °C. Sample RCA5 was collected at 62.4 °C from a separate slow-moving outflow of Smoking Gun Spring, ∼ 20 m from the source. pH values across all RCA samples were similar (9.1-9.4). RCA5 had the highest sulfide concentration (18.99 µM) compared to the other RCA samples which ranged from 0.25-0.44 µM (Table 1; Supplementary File 1).

In the Lower Geyser Basin, Sentinel Meadows (SM) samples were collected from shallow outflows of three separate springs, all close to their source (5-12 m downstream). Samples SM1, SM2 and SM4 all had pH near 8, while temperatures were 68.4 °C, 51.4 °C, and 62.3 °C, respectively (Table 1). White Creek Area (WCA) samples were among the highest in temperatures (69.4-73 °C) and consisted of exclusively green filaments (Table 1). Beyond community morphologies and temperatures, other distinguishing physical and geochemical parameters were not observed for WCA samples, although samples 18WCA1 and 18WCA2 were the only samples for which nitrate and ammonium concentration data are available. Nitrate was detectable in both samples (14.3 µM), while ammonium was below detection limit in 18WCA1 and 6.0 µM in 18WCA2 (Supplementary File 1). Micromolar concentrations of nitrate and/or ammonia have been previously reported LGB springs (Havig et al., 2021; Holloway et al., 2011), and thus we expect 2017 samples would have had similar concentrations. It is possible that unmeasured parameters and/or biogeography are primary predictors of the characteristic extremely thermophilic green filaments in these springs. Putative A’-like cyanobacterial MAGs were dominant among these samples, followed by *Chloroflexus* and *Roseiflexus* (Fig 2). Imperial Geyser Basin samples include BG1and IGB1. The BG1 sample, at 8.7 pH and 68.5 °C, consisted of a red mat with filaments, while IGB1 was taken from an orange/yellow mat at pH 8.2 and 66.6 °C. Samples in the Lower Geyser Basin had sulfide concentrations ranging from 0.22-4.93 µM.

**Figure 2.**
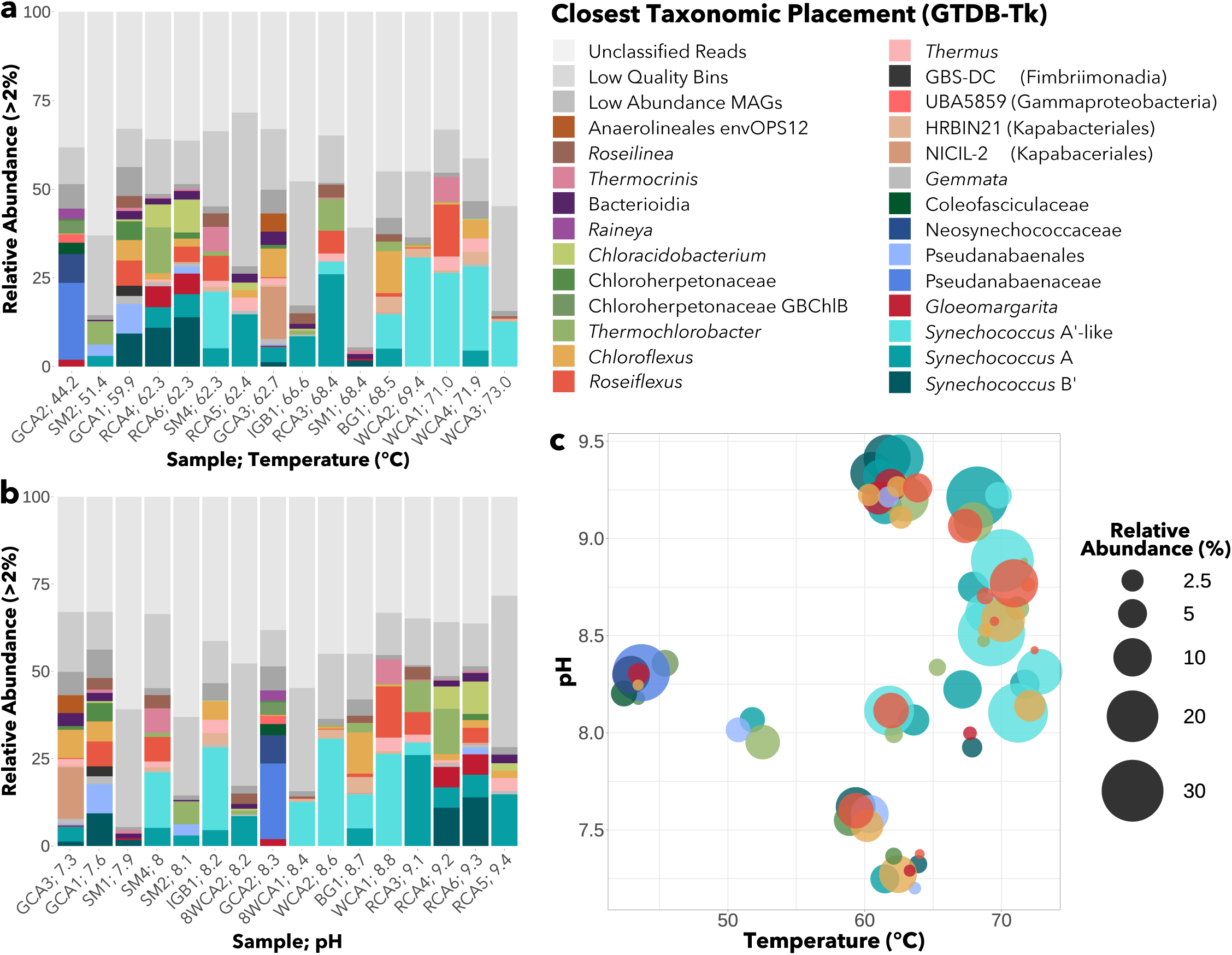
MAG abundance across samples. Relative abundance of all MAGs with > 2% abundance in at least one sample. Samples are ordered by **(a)** temperature or **(b)** pH. **(c) A**bundances of phototrophic MAGs along combined pH and temperatures. Unmapped reads, low quality bins, and low abundance MAGs are grouped per sample (shown in grey).

In the Gibbon Geyser Basin, samples taken from the Geyser Creek Area (GCA) consisted of primarily of orange, pink, and red mats overlaid on black and grey sediments. Two samples (GCA1 and GCA2) were from a single shallow outflow within ∼10 m of the source spring, while a third (GCA3) was from a shallow outflow further downstream of its source. Temperatures in the GCA ranged from 44.2-62.7°C and pH from 7.3 to 8.3 (Table 1). Sulfide concentrations ranged from 0.62-2 µM while sulfate (1.2-1.4 mM) and chloride (11-13 mM) were elevated compared to other sample areas (∼0.2 mM and 6-9 mM respectively) (Supplementary File 1).

A total of 360 metagenome bins were recovered in this study, 221 of which met minimum criteria to be considered MAGs (> 50% completeness, < 10% contamination). Of the 221 MAGs recovered, 175 MAGs were considered high-quality (> 70% completeness). Taxonomic abundance estimates of MAGs revealed that Cyanobacteria, Chloroflexi, and Chlorobi were the most common photosynthetic phyla observed across samples (Fig 2) consistent with previous work in YNP hot springs (Hamilton et al., 2019; reviewed in Tank et al., 2017).

### Recovery of novel Synechococcus-like A/B-lineage MAGs from the upper temperature limit of photosynthesis

Our primary focus in this study is with *Synechococcus* A/B-lineage (genus-level) MAGs, as they were present across nearly all sites (Fig. 2), and the majority – 22 MAGs – had > 50% completeness and < 10% contamination, with a high-quality subset of 15 MAGs at > 70% completeness. A/B-lineage MAGs were recovered from all samples between 51.4°C and 73°C, with three distinct taxa identified by GTDB-Tk (Chaumeil et al., 2019). Two other MAGs classified at the order and family level with the A/B-lineage were recovered in low relative abundance from the two lowest temperature samples (GCA2.22 = 1.13% at 44.2°C; SM2.3A = 0.57% at 51.4°C).

Of the 22 total A/B-lineage MAGs (> 50% completeness), ten had greater than 99% average nucleotide identity (ANI) with the closest reference genome from strain OS-A. Meanwhile five MAGs had ∼96-98% ANI with OS-B’ – suggesting relatively higher diversification among B’ MAGs recovered (Supplementary File 2). MAGs assigned to the A taxon were recovered from temperatures between 51.4 and 71.9°C, while B’ MAGs occupied temperatures between 59.9 and 68.4°C but were most abundant below 62.3°C (Fig 2B, Fig 3). The remaining seven A/B-lineage MAGs recovered from 62.3-73.0°C samples had ∼85-88 % identity to the OS-A reference genome. MAGs representing this taxon were most abundant at the highest temperatures in our samples (69-73°C; Fig 2B), suggesting they are A’-like ecotypes (Ferris & Ward, 1997; Ward et al., 2006). The two MAGs classified near the A/B-lineage at the order or family level had no closely related reference genome available. A phylogenomic analysis using 71 concatenated marker genes from strictly high-quality (> 70% completeness) cyanobacterial MAGs (26 total in this study) and reference genomes showed that these two MAGs each formed their own clade, while confirming tight clustering of the three A/B-lineage clades (Fig S1).

**Figure 3.**
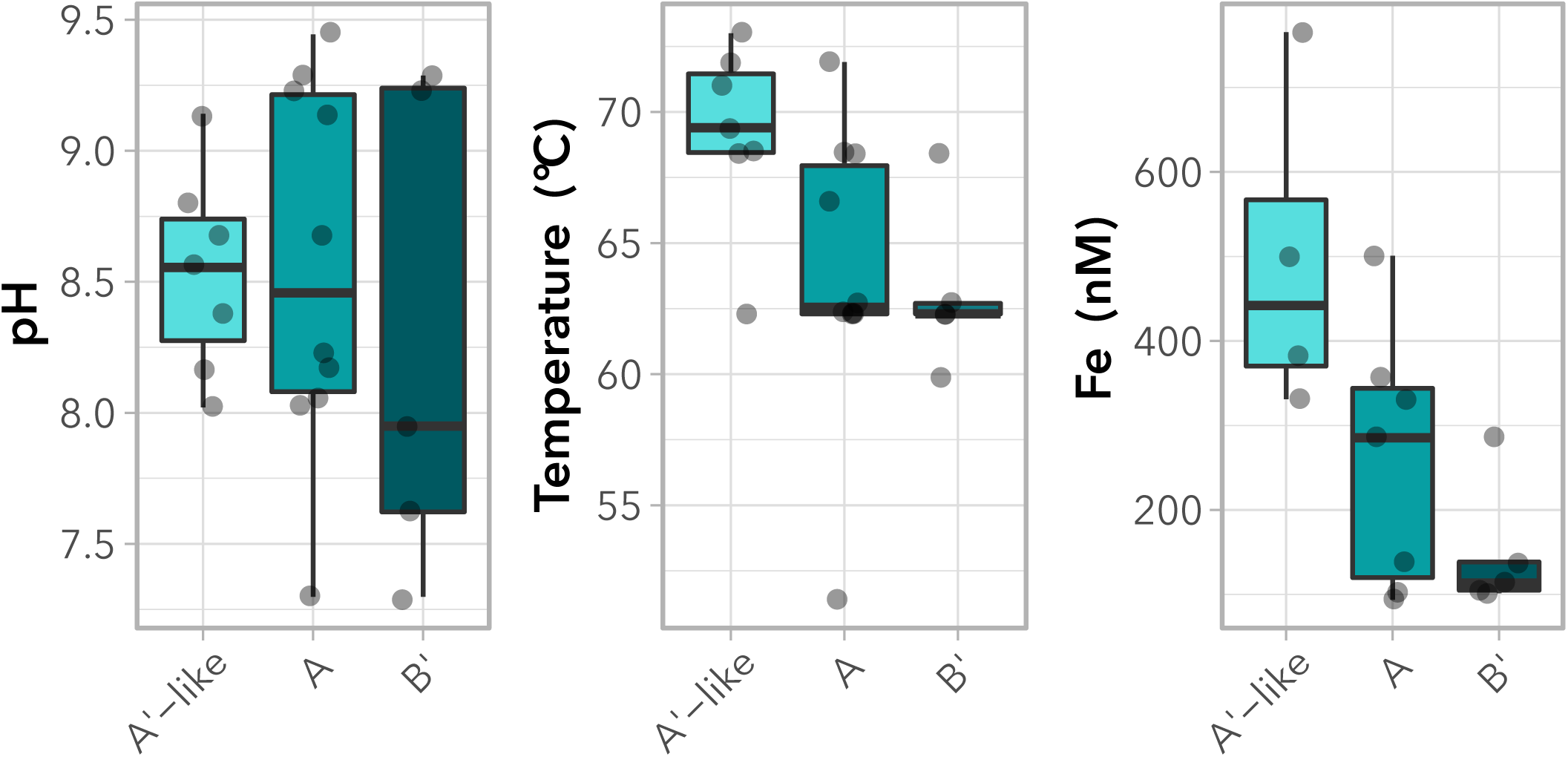
*Synechococcus*-like A/B-lineage MAG ranges with environmental parameters. Presence of cyanobacterial MAGs by taxon across pH, temperature, and Fe concentrations.

To better define the taxonomy of the suspected A’-like clade of MAGs, we performed a phylogenetic analysis of PsaA and RbcL sequences recovered from 15 high-quality A/B-lineage MAGs (Table 2). PsaA sequences recovered from our putative A’-like MAGs cluster with sequence PEA9 from another putative A’-like ecotype in YNP (Becraft et al., 2011) (Fig S2). RbcL sequences recovered from A’-like MAGs cluster with a Synechococcus-like isolate, strain OH28, which has also been classified as an A’-like ecotype (Miller et al., 2013). These observations suggest that this highest temperature clade of *Synechococcus*-like A/B-lineage MAGs most likely constitutes a A’-like ecotype, for which no genomes had been previously sequenced.

**Table 2.** Cyanobacterial MAG descriptions. RCA=Rabbit Creek Area ; SM = Sentinel Meadows ; BG = Boulder Geyser ; WCA = White Creek Area ; GCA = Geyser Creek Area ; IGB = Imperial Geyser Basin

| MAG | Relative Abund (%) | Completeness | Contamination | Taxonomic Assignment | Closest Reference Genome | ANI to Ref. (%) | High Qual.? |
| --- | --- | --- | --- | --- | --- | --- | --- |
| RCA3.9B | 3.51 | 64.47 | 0.52 | *Synechococcus*-like A' type | JA-3-3Ab [OS-A] | 85.16 |  |
| BG1.9A | 9.69 | 79.82 | 4.87 | *Synechococcus*-like A' type | JA-3-3Ab [OS-A] | 87.29 | ✓ |
| 18WCA2.12A | 23.65 | 81.58 | 4.39 | *Synechococcus*-like A' type | JA-3-3Ab [OS-A] | 87.32 | ✓ |
| WCA2.4 | 30.76 | 86.84 | 0 | *Synechococcus*-like A' type | JA-3-3Ab [OS-A] | 87.3 | ✓ |
| SM4.3A | 15.86 | 94.74 | 0.88 | *Synechococcus*-like A' type | JA-3-3Ab [OS-A] | 87.11 | ✓ |
| WCA1.12 | 26.42 | 96.49 | 0 | *Synechococcus*-like A' type | JA-3-3Ab [OS-A] | 87.47 | ✓ |
| 18WCA1.11 | 12.68 | 98.25 | 0 | *Synechococcus*-like A' type | JA-3-3Ab [OS-A] | 87.47 | ✓ |
| 18WCA2.12B | 4.48 | 51.27 | 0 | *Synechococcus*-like A type | JA-3-3Ab [OS-A] | 99.61 |  |
| RCA6.5B | 6.51 | 55.78 | 0.44 | *Synechococcus*-like A type | JA-3-3Ab [OS-A] | 99.32 |  |
| BG1.9B | 4.99 | 62.96 | 0 | *Synechococcus*-like A type | JA-3-3Ab [OS-A] | 99.24 |  |
| SM4.3B | 5.14 | 64.04 | 0 | *Synechococcus*-like A type | JA-3-3Ab [OS-A] | 99.21 |  |
| RCA4.11B | 5.81 | 72.37 | 0 | *Synechococcus*-like A type | JA-3-3Ab [OS-A] | 99.21 | ✓ |
| SM2.9A | 3.00 | 76.41 | 4.66 | *Synechococcus*-like A type | JA-3-3Ab [OS-A] | 99.31 | ✓ |
| RCA3.9A | 25.99 | 86.55 | 0.88 | *Synechococcus*-like A type | JA-3-3Ab [OS-A] | 99.15 | ✓ |
| GCA3.42A | 4.28 | 89.3 | 0.1 | *Synechococcus*-like A type | JA-3-3Ab [OS-A] | 99.23 | ✓ |
| IGB1.12 | 8.54 | 91.23 | 0.1 | *Synechococcus*-like A type | JA-3-3Ab [OS-A] | 99.34 | ✓ |
| RCA5.7 | 14.73 | 94.3 | 0.88 | *Synechococcus*-like A type | JA-3-3Ab [OS-A] | 99.16 | ✓ |
| RCA6.5A | 13.88 | 53.9 | 4.39 | *Synechococcus*-like B' type | NA | NA |  |
| RCA4.11A | 10.92 | 59.91 | 3.2 | *Synechococcus*-like B' type | JA-2-3B'(2-13) [OS-B'] | 96.96 |  |
| SM1.7B | 1.66 | 71.93 | 8.33 | *Synechococcus*-like B' type | JA-2-3B'(2-13) [OS-B'] | 97.99 | ✓ |
| GCA3.42B | 1.22 | 72.19 | 0 | *Synechococcus*-like B' type | JA-2-3B'(2-13) [OS-B'] | 98.12 | ✓ |
| GCA1.24 | 9.32 | 91.23 | 1.75 | *Synechococcus*-like B' type | JA-2-3B'(2-13) [OS-B'] | 97.93 | ✓ |
| GCA2.22 | 1.13 | 98.72 | 0 | A/B lineage Family | NA | NA | ✓ |
| SM2.3A | 0.57 | 82.32 | 7.42 | A/B-lineage Order | NA | NA | ✓ |
| RCA6.18 | 5.85 | 62.73 | 3.42 | *Gloeomargarita* | *G. lithophora* Achichica-D10 | 79.93 |  |
| GCA2.30 | 1.94 | 64.14 | 0 | *Gloeomargarita* | *G. lithophora* Achichica-D10 | 83.51 |  |
| RCA4.1 | 5.88 | 64.67 | 1.71 | *Gloeomargarita* | *G. lithophora* Achichica-D10 | 79.91 |  |
| GCA3.22 | 0.39 | 84.62 | 0 | *Gloeomargarita* | *G. lithophora* Achichica-D10 | 79.43 | ✓ |
| SM1.2 | 0.56 | 86.05 | 2.32 | *Gloeomargarita* | *G. lithophora* Achichica-D10 | 79.2 | ✓ |
| GCA1.18 | 0.53 | 74.21 | 0.59 | Pseudanabaenaceae bacterium | Pseudanabaenaceae genus M5B4 | 77.84 | ✓ |
| GCA2.3 | 21.65 | 88.09 | 0.24 | Pseudanabaenaceae bacterium | Pseudanabaenaceae genus M5B4 | 78.63 | ✓ |
| SM2.6A | 0.68 | 89.27 | 1.65 | Pseudanabaenales bacterium | NA | NA | ✓ |
| SM2.6B | 2.48 | 74.4 | 5.07 | Pseudanabaenales bacterium | NA | NA | ✓ |
| RCA6.13 | 1.89 | 91.04 | 0.47 | Pseudanabaenales bacterium | NA | NA | ✓ |
| GCA3.6 | 0.43 | 95.28 | 0.63 | Pseudanabaenales bacterium | NA | NA | ✓ |
| GCA1.15 | 7.77 | 95.4 | 2.67 | Pseudanabaenales bacterium | NA | NA | ✓ |
| GCA2.36 | 3.22 | 94.52 | 0.67 | Coleofasciculaceae bacterium | Coleofasciculaceae bacterium JAAUUE01 | 75.88 | ✓ |
| GCA2.6 | 8.12 | 85.26 | 0.24 | Neosynechococcaceae bacterium | Neosynechococcaceae GCF-001939115 | 75.91 | ✓ |

### Recovery of Psuedanabaenales MAGs and novel thermophilic Gloeomargarita MAGs

Two other sets of cyanobacterial MAGs that we recovered are those belonging to Pseudanabaenales, and *Gloeomargarita* taxa. A total of seven Pseudanabaenales MAGs were recovered from samples up to 62.7 °C and pH 7.3-9.3 in the Geyser Creek Area, Rabbit Creek Area, and Sentinel Meadows. Two MAGs placed within the family *Pseudanabaenaceae* – GCA1.18 and GCA2.3 – had ∼78% ANI to cyanobacterial MAG M5B4 recovered from a hot spring in Lakash India (Roy et al., 2018) (Fig. S1, Table 2). Four other MAGs – GCA3.6, GCA1.15, SM2.6, and RCA6.13 – had unknown taxonomic placement within the order Pseudanabaenales. A seventh MAG, SM2.6B, was originally assigned to the *Pseudanabaenaceae* family by GTDB-Tk. However, a phylogenomic analysis using 71 concatenated bacterial marker genes (Fig S1) from 26 high quality cyanobacterial MAGs (>70% completion, <10% contamination) placed SM6.2B nearer to the clade of four unknown Pseudanabaenales MAGs.

A total of seven MAGs (2 high-quality) were assigned to genus *Gloeomargarita*, with∼79-82% ANI to the closest reference genome *G. lithophora* Alchichica-D10 (Couradeau et al., 2012; Moreira et al., 2017). Phylogenomic analysis showed that two high-quality *Gloeomargarita* MAGs form a single tight clade associated with Alchichica-D10 (Fig S1). *Gloeomargarita* MAGs were recovered from samples with temperatures up to 68.4°C and from samples between pH 7.3 and 9.3 (Fig 2), but were most abundant (> 2%) at pH 9.2-9.3. This distribution is consistent with previous 16S rRNA amplicon results indicating the presence of *Gloeomargarita* OTUs up to 70.8°C in Rabbit Creek and Bison Pool (Bennett et al., 2020). In comparison, the *G. lithophora* Alchichica-D10 isolate has a growth range of 15-30°C in the lab (Couradeau et al., 2012; Moreira et al., 2017). Considering *Gloeomargarita* is an early branching genus of cyanobacteria (Couradeau et al., 2012; Moreira et al., 2017; Ponce-Toledo et al., 2017), recovery of MAGs (and potentially future isolates) at such high temperatures, may yield new insights into the evolution of thermophily in cyanobacteria.

### Influence of environmental parameters on A’-like MAG distributions

Temperature, pH, and sulfide have all been demonstrated to be the primary environmental constraints on photosynthesis in hot springs (Boyd et al., 2012; Cox et al., 2011; Hamilton et al., 2012). However, in our samples, the only environmental parameters that impacted the distribution of A’-like vs B’-like MAGs were temperature (P=0.0249) and Fe concentrations (P=0.0149; Dunn’ s test with Bonferroni correction) (Fig 3). A’-like MAGs were generally recovered from outflows with higher temperatures (62-73 °C) than B’ MAGs (60-68 °C). Differences in temperature ranges between A’-like and B’ MAGs are consistent with previous work showing shifted, yet overlapping, temperature ranges among B’, A, and A’ ecotypes, both *in situ* and in laboratory cultures (Allewalt et al., 2006; Ferris & Ward, 1997; Miller & Castenholz, 2000). A’-like MAGs were also recovered from samples with higher iron concentrations (383-501 nM Fe) than B’ MAGs (101-285 nM Fe). It is possible that the impacts of temperature and iron on the distribution of A’-like MAGs are interlinked. Phototrophs require high levels compared to other organisms (Kranzler et al., 2013) because it is essential for the construction of photosystem II (PS II), chlorophyll synthesis, and is an essential cofactor for photosystem I (PS I) and the photosynthetic electron transport chain. A higher iron concentration in samples where A’-like MAGs were recovered compared to other samples in the study might suggest higher demands for PS repair and turnover in cyanobacteria near the upper temperature limit, potentially due to increased heat stress on the photosynthetic apparatus.

Beyond differences in iron concentration and temperature, A’-like MAGs were recovered primarily from filaments. These include four green filament samples (WCA1, WCA2, 18WCA1, 18WCA2), and one red mat/filaments sample (BG1). A’-like MAGs were also recovered from two of orange mats (RCA3 and SM4). B’ MAGs were only recovered from mat communities, while A MAGs were recovered from eight mat communities and two filament communities. Given that green filament samples containing A’-like MAGs were only observed in White Creek Area samples, we expected biogeography to impact phylogenomic distance among A’-like, A and B’ clades. An influence of biogeography on variation in *psaA* genes among A/B-lineage organisms has been previously demonstrated (Becraft et al., 2020). Accordingly, geographic distance was significantly predictive of phylogenomic distance among the A/B-lineage (Mantel test; P=0.014, r = 0.282). Although the Pearson’s r coefficient indicates weak correlation, this result suggests that dispersal barriers are a factor in the distribution of A/B-lineage MAGs. Thus, temperature, iron concentration, and biogeography all likely play roles in the distribution of A’-like MAGs, and the community morphology observed in WCA samples.

A’-like MAGs were only recovered from pH of 8.0-9.1 compared to A (7.3-9.4) and B’ MAGs (7.3-9.3) (Fig 2B; Fig 3), suggesting that one or multiple physiological components have tighter pH constraints in A’-like MAGs, perhaps as a consequence of adaptation to the highest temperatures at which phototrophy has been observed. In general, while sample size is limited, all phototrophic fringe communities observed above 70°C in this study were limited to pH 8.1-8.8 (Fig 2C). Similar constraints on phototrophs observed above 70°C were previously seen across much larger sample sets within YNP (Hamilton et al., 2012, 2019). In those studies, *chlL*/*bchL* and *bchY* amplicons at sample temperatures between 65°C and 70°C were observed between pH 5.5 and pH 9.5, while amplicons above 70°C were more narrowly limited to pH ∼7.5-9.2.

### Pangenomes of A/B-lineage metagenome-assembled and reference genomes

Given that A/B-lineage taxa were the most widely distributed among cyanobacterial MAGs we recovered, and that each phylogenomically distinct clade occupies differing temperature ranges, we narrowed our focus to A/B-like MAGs to elucidate shared and unique genome content among them. We performed pangenome analysis within anvi’o (Eren et al., 2020) using 15 strictly high quality (> 70% completeness) A/B-lineage *Synechococcus*-like MAGs and the OS-A and OS-B’ reference genomes. MAGs with completeness below 70% were excluded to decrease the number of missing genes due to metagenome assembly and binning error. The pangenome (Fig 4) was composed of a total of 3274 gene clusters, with a core genome of 489 universally shared clusters. An auxiliary core of 1229 gene clusters was shared by multiple representatives in each of the A’-like, A, and B’ clades. Of particular interest were gene clusters that were unique to or absent from A’-like MAGs, given that they are capable of occupying the thermal limit of photosynthesis and that they form a separate clade from A and B’ genomes (Fig S1).

**Figure 4.**
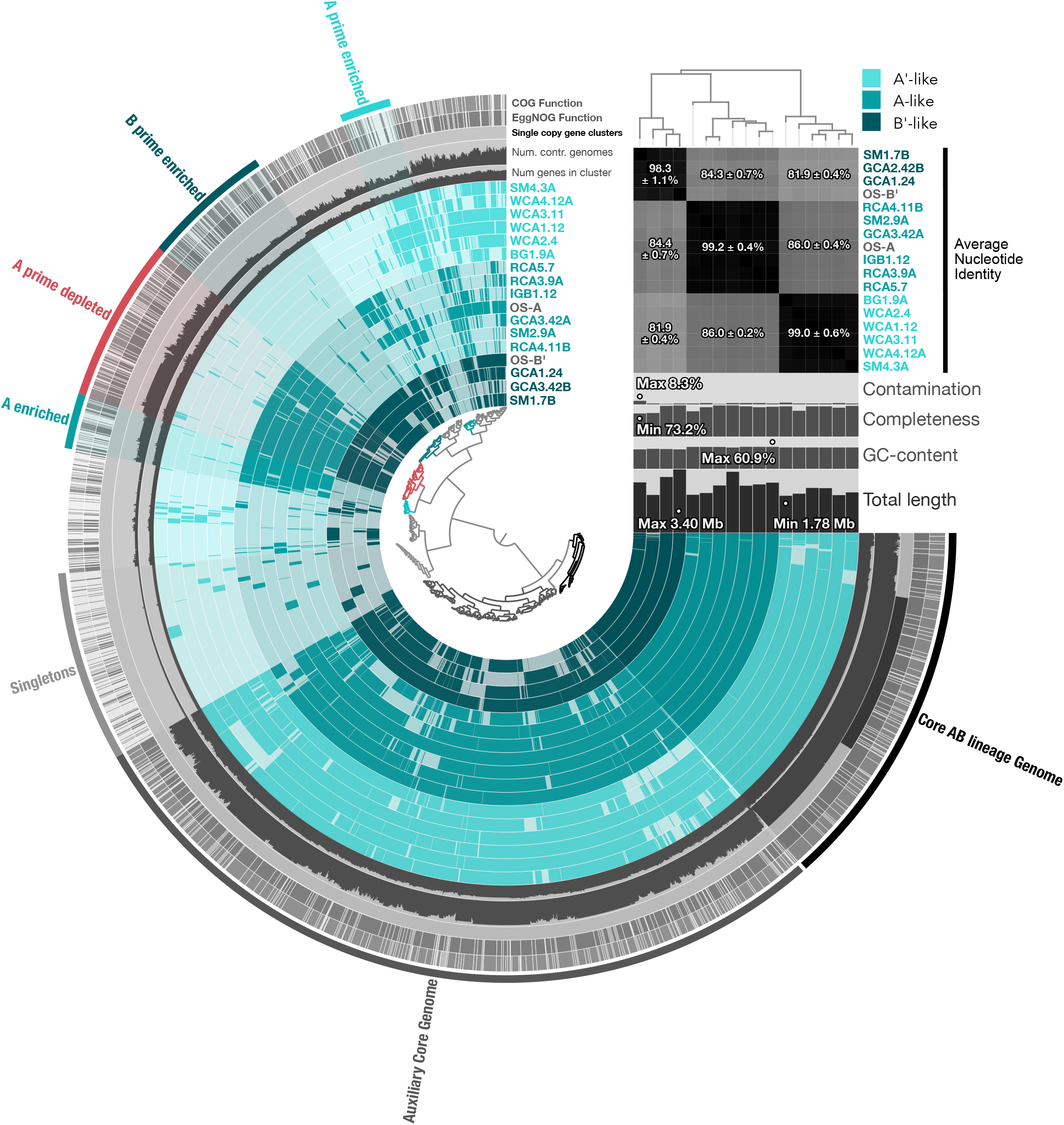
Pangenome of *Synechococcus*-like A/B lineage MAGs. Gene clusters present in 17 high quality (> 70% completion, < 10% contamination) A/B-lineage MAGs are shown in colored rings. Central dendrogram depicts ordering of gene clusters by frequency (D: Euclidian, L: Ward), and was used to select gene clusters unique to, or enriched in different MAGs or clades (outermost rings and highlighted gene cluster selections). Percent average nucleotide identity comparisons are statistically represented on a clade-by-clade basis. Contamination and completion were calculated by checkM and the anvi’o pangenomics suite and ANI was calculated by pyANI within anvi’o. NCBI clusters of orthologous groups (COGs) were identified using the 2020 release within anvi’o.

### Recovered A’-like MAGs do not encode nitrogenase and DPOR

To examine potential gene gain or loss within A’-like strains that may suggest adaptation to higher temperatures, a functional enrichment analysis was performed within anvi’ o (Shaiber et al., 2020). Here, the term ‘enriched’ is used when a gene cluster or a set of gene clusters is present more often in one clade of MAGs (A’-like, A or B’) than the others and supported by an adjusted q-value < 0.05 in the functional enrichment analysis. Likewise, a gene cluster is considered ‘depleted’ in a clade when found more often in two others. Several gene clusters were depleted (258 clusters), rather than enriched (80 clusters), in the A’-like clade, which suggests a streamlining of the A’-like genome to accommodate a high-temperature range. These results are consistent with a general trend previously noted for thermophilic cyanobacteria including the A/B lineage (Alcorta et al., 2020; Larsson et al., 2011). Gene clusters depleted in A’-like MAGs include those predicted to encode nitrogenase — *nifH, nifD, and nifK* (Supplementary File 3). Although functional enrichment scores for individual *nif* genes were just above the 0.05 threshold to be considered statistically significant (q=0.0514), a consistent qualitative pattern emerges in the representation of *nif* operons and accessory genes across MAGs (Fig 5, Table S2). OS-A and OS-B’ reference genomes and at least two MAGs assigned to each of these clades encode a nitrogenase enzyme while these genes were absent from A’-like clade. Other genes necessary for nitrogenase assembly are similarly absent from A’-like MAGs.

**Fig 5.**
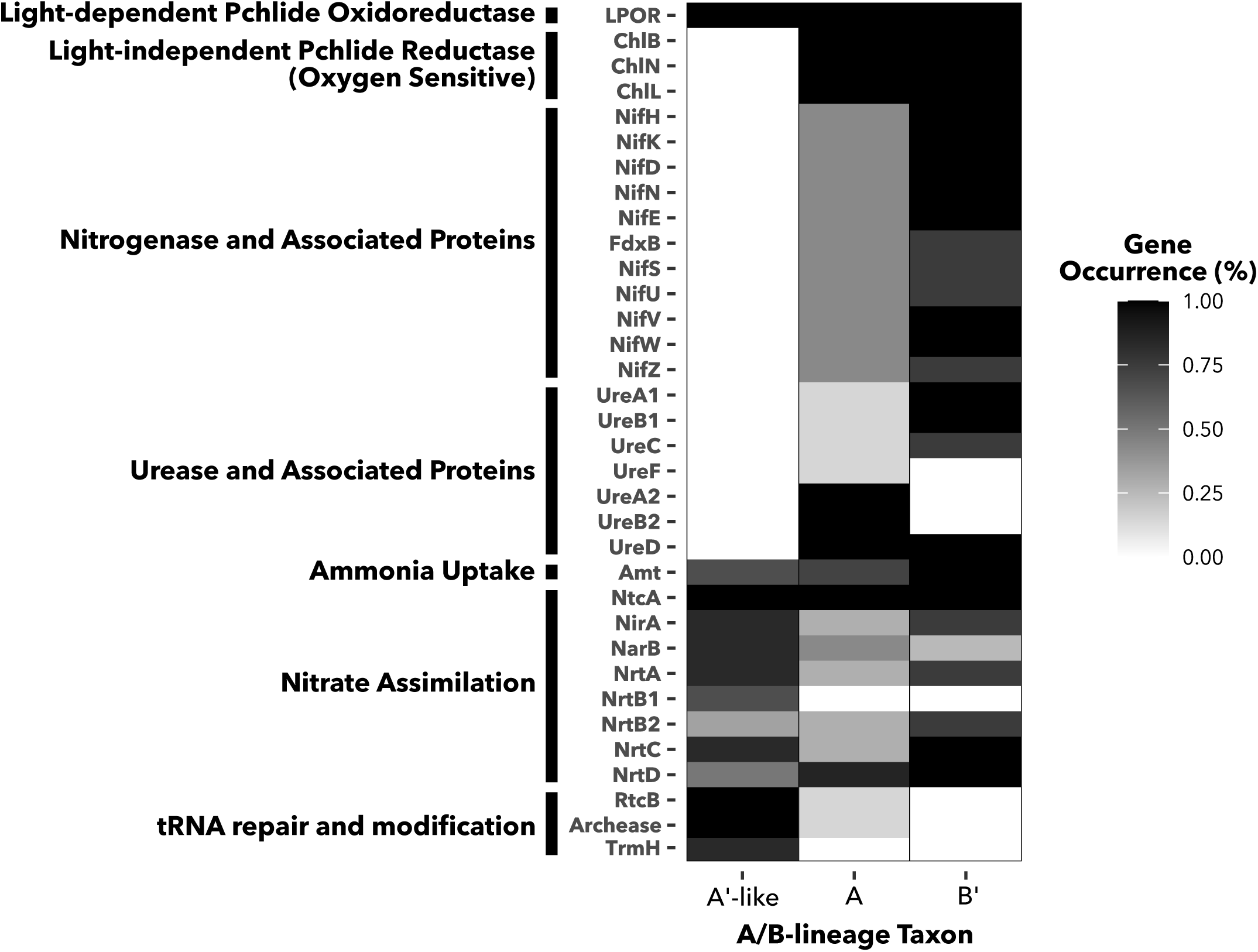
Occurrence of chlorophyllide synthesis, nitrogen metabolism, and tRNA repair and modification genes in A/B-lineage taxa. Heatmap fill indicates percentage of A’-like (N=6), A (N=7) and B’ (N=4) MAGs in which each gene cluster is present. Gene clusters are labeled by names of predicted proteins they encode and ordered by function.

These results suggest that A’-like cyanobacteria adapted to the upper temperature limits of photosynthesis must rely on fixed nitrogen. Unlike A and B’ MAGs and reference genomes, which encode clade-specific variants of urease (Bhaya et al., 2007), no urease genes were recovered from A’-like MAGs (Fig 5). The ammonium transporter Amt along with a full suite of nitrate assimilation genes were present in the A’-like clade, suggesting that nitrate and/or ammonia could be the primary nitrogen sources for these organisms. As a likely source of fixed nitrogen, genes encoding nitrogenase were present in all metagenomes where A’-like MAGs were recovered except the highest temperature sample (18WCA1, 73°C). In 18WCA1, the only nitrogenase genes present in the metagenome assembly were a *nifHBDK* cassette from *Roseiflexus*. No associated maturation genes were present, consistent with the genomes of *Roseiflexus* isolates, for which no evidence of diazotrophy has been observed (Hanada et al., 2002; van der Meer et al., 2010). However, micromolar nitrate detected in 18WCA1 suggests the overflowing water may provide bioavailable nitrogen. Thus, we expect the overflowing water and/or diazotrophy by other members of the community to be the most likely sources of fixed nitrogen for A’-like MAGs.

The genes *chlLNB*, which encode the dark-operative protochlorophyllide reductase (DPOR), an oxygen-sensitive enzyme sharing evolutionary history with nitrogenase, were also absent in A’-like MAGs. Conversely, *chlLNB* were present in all A type and B’ type MAGs and in the OS-A and OS-B’ reference genomes. In the absence of DPOR, the oxygen-tolerant light-dependent protochlorophyllide oxidoreductase (LPOR) is essential for chlorophyllide synthesis. Accordingly, the gene encoding LPOR, *por*, was found in all A/B-lineage MAGs and reference genomes, including A’-like MAGs. We can only speculate on the mechanisms that drove the A’-like clade to lose both nitrogenase and DPOR. Given the oxygen-sensitive nature of nitrogenase and DPOR, a loss of both enzymes in A’-like MAGs might suggest a loss of the ability to protect them. However, with the short distance of these samples from the source spring, low solubility of oxygen at these high temperatures (68-73°C), and expression of both of these enzymes at night when mats are anoxic (Steunou et al., 2006, 2008). we do not expect oxygen sensitivity to be an evolutionary force driving their loss. Instead, the availability of nitrate/ammonium from overflowing water, and the presence of genes for diazotrophy and LPOR in the metagenomes, we expect loss of nitrogenase and DPOR to be most likely due to their dispensability. The evolutionary history of diazotrophy is replete with examples of intra- and inter-phylum gene transfers as well as trait loss (Hartmann & Barnum, 2010; Latysheva et al., 2012; Raymond et al., 2004). Similar trait loss of DPOR is common to many photosynthetic taxa, often associated with maintenance of multiple *por* genes (Hunsperger et al., 2015; Vedalankar & Tripathy, 2018). Loss of these genes in the highest temperature clade of A/B-lineage MAGs is also consistent with a general trend toward genome reduction in thermophiles (Alcorta et al., 2020; Larsson et al., 2011),

### The A’-like clade of MAGs is enriched for non-cyanobacterial tRNA damage response proteins

Two gene clusters enriched in the A’-like clade of the A/B-lineage pangenome are those encoding a tRNA-splicing ligase, RtcB, a predicted archease involved in RtcB activation, and a tRNA methyltransferase, trmH (Supplementary File 3). Archaease and RtcB are conserved across all domains of life (Desai et al., 2014) and are thought to assist in recovery from stress-induced tRNA damage (Santamaría-Gómez et al., 2021; Tanaka & Shuman, 2011), while TrmH is implicated in regulating transcriptional response to stress conditions in *E. coli* (Galvanin et al., 2020). Archease sequences were only recovered from A’-like MAGs and the OS-A genome. A BLASTP search of A’ RtcB against the NCBI nr database returned hits belonging to a small group of A/B-lineage genomes (∼96% identity over 100% query coverage), including the OS-A genome, followed by RtcB sequences belonging to members in the Methylothermaceae family (∼75-76 % identity over 100% query coverage). BLASTP searches of A’-like archease against the full nr database produced similar results. These results suggest RtcB and archease in the A/B lineage are highly diverged from those in other cyanobacteria. The top hit resulting from a BLASTP search for A’-like RtcB against all other cyanobacteria was to a tRNA-splicing ligase in a *Cyanobium* species at ∼60% identity over 100% query coverage, while other hits were below 50% identity. Furthermore, the top hit for A’-like archease against cyanobacteria outside the A/B-lineage only had ∼30% identity over 99% gene coverage. These results indicate that archease and RtcB in the A/B-lineage diverged from other cyanobacteria early in their history. Similarly, top hits from A’-like TrmH sequences searched against the NCBI nr database were from Firmicutes, Proteobacteria, Actinobacteria, Chloroflexi, and the Fibrobacters/Chlorobi/Bacteriodetes (FBC) superphylum. Meanwhile, no significant similarity was found in BLASTP searches of A’-like TrmH against strains OS-A and OS-B’. Altogether, these results suggest a difference in tRNA repair and modification in response to stress between A’-like organisms and other A/B lineages, perhaps acquired through horizontal gene transfer.

### Functional divergence of conserved genes in Synechococcus-like MAGs

We asked whether any genes present in A’-like MAGs diverged from genomes in the A and B’ clades in ways that may indicate adaptation to higher temperatures. Among gene clusters that were conserved across clades, we queried those without paralogs, with functional homogeneity (defined by point mutations) below 85% and geometric homogeneity (defined by gaps) above 85%. We compared phylogenetic distance matrices of the 22 annotated gene clusters that met these criteria to distance matrices produced from a phylogenomic tree of 71 conserved marker genes in A/B-lineage MAGs (Fig 6A). We then sought gene clusters with a pattern of divergence that places A’-like sequences further away from A and B’ sequences than would be predicted by the phylogenomic tree – a pattern that would be suggestive of functional adaptation in the A’-like clade. One such cluster encodes a transcription factor in the sigma-70 family, which displays a clear pattern of divergence of A’-like sequences away from A and B’ sequences (Fig 6B). Although this protein is uncharacterized, its divergence from other A/B-lineage types represents a potential regulon that is specialized in A’-like organisms.

**Figure 6.**
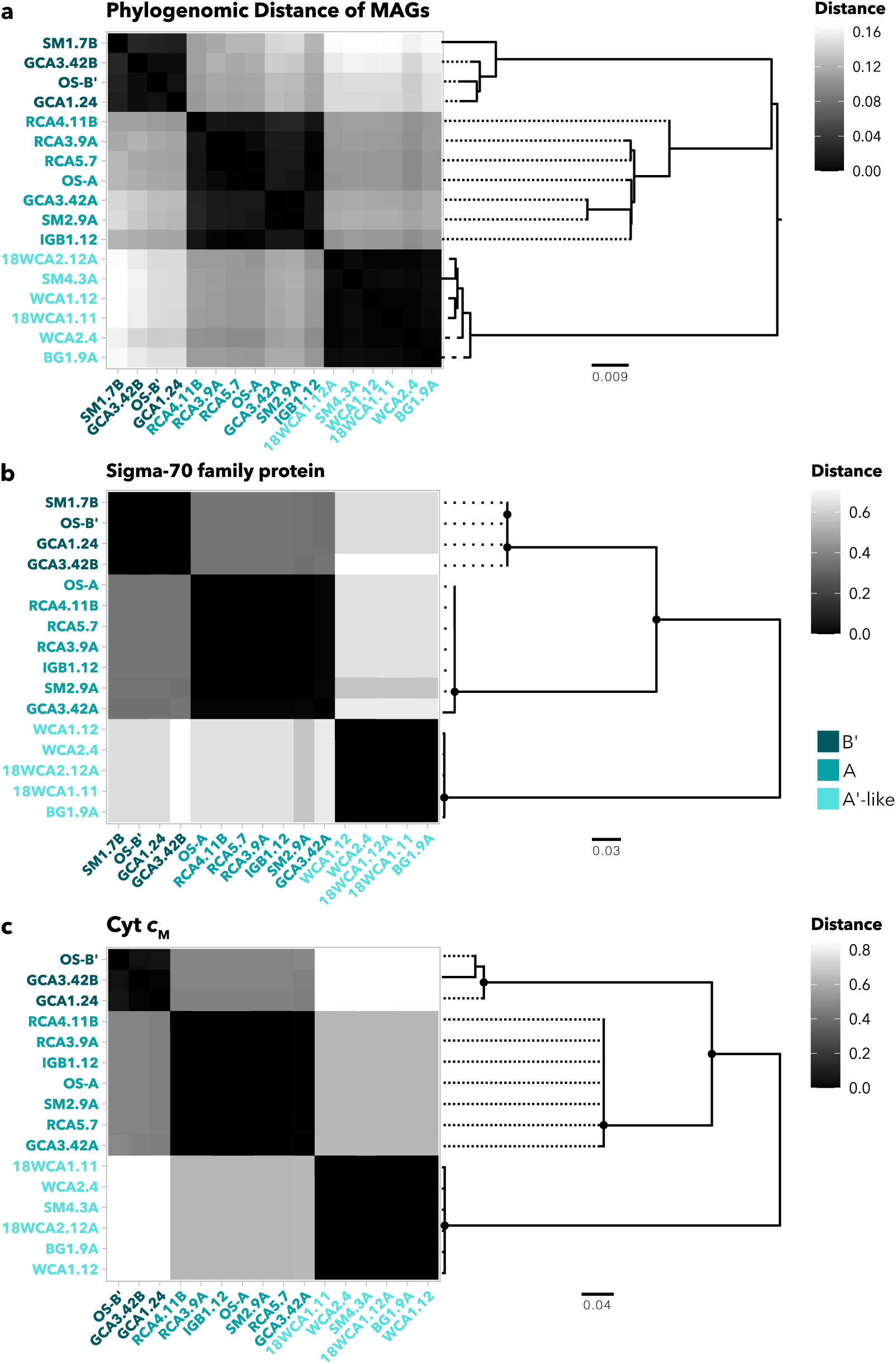
Phylogenetic distances of select gene clusters. **(a)** Phylogenomic distance heatmap and corresponding tree using an aligned, concatenated set of 71 conserved marker genes from A/B-lineage MAGs, OS-A, and OS-B’ reference genomes. **(b)** Phylogenetic distances of *cytM* genes extracted from A/B-lineage MAGs. **(c)** Uncharacterized sigma-70 family protein extracted from A/B-lineage MAGs and reference genomes. Node symbols denote bootstrap values above 80 from 100 bootstraps within RAxML. Each MAG is colored according to its taxon (A’-like, A, and B’).

Another gene cluster with high phylogenetic distance between the A’-like clade and the other two A/B-lineage clades encodes cytochrome *c*_M_ (Fig 6C), which is present across cyanobacterial taxa, but whose physiological role is still debated (Bialek et al., 2016; Solymosi et al., 2020). Transcription of *cytM* increases under conditions such as low temperature, high intensity light, oxidative stress, osmotic stress, and nitrogen and sulfur limitation (Ludwig & Bryant, 2012a, 2012b). It has been suggested that under these stress conditions, Cyt *c*_M_ either plays a regulatory role, or it alleviates photosynthetic over-reduction of electron carriers (Bernroitner et al., 2009; Bialek et al., 2016; Hiraide et al., 2015). Furthermore, studies in *Synechocystis* 6803 have shown that Cyt *c*_M_ suppresses both heterotrophy in the dark and photosynthesis during photomixotrophy (Hiraide et al., 2015; Solymosi et al., 2020), supporting the notion that it performs a regulatory function. Divergence in Cyt *c*_M_ between A’-like MAGs and other A/B-lineage genomes may suggest they differ in their response to stress conditions. Since A’-like populations often occupy the absolute limits of photosynthesis, it is possible that temperature fluctuations require faster or stronger suppression of photosynthesis. Alternatively, since *cytM* is induced under N-limitation, it is possible that a reliance on bioavailable nitrogen from overlying water or neighboring microbes may have required modification of the Cyt *c*_M_ regulon. Similar to phylogenomic distances, a clear influence of biogeography on intra-clade variation of sigma-70 factor or Cyt *c*_M_ was not observed, particularly among sequences from the A’-like clade. Since intraclade variation was small, we conclude that variation observed is clade-specific. Considering their potential to affect responses to environmental fluctuations, divergence in both Cyt *c*_M_ and this sigma-70 factor, may guide future transcriptomic work into differences in stress responses between A/B-lineage clades in YNP hot springs. For a full summary of gene clusters, including annotations, functional / geometric homogeneity scores and amino acid sequences, see Supplementary File 4.

## Conclusions

Cyanobacteria in Yellowstone hot springs have been widely studied for decades, yet much of the work has focused on thick laminated mats in only two hot springs within the same geyser basin. Here, we expanded this large body of work to include metagenome assembled genomes recovered from hot springs spanning different regions of YNP, with varying temperatures, pH, geochemistry and community morphologies. Particularly, our work focused on genomic divergence among three clades or ecotypes (A, B’, A’-like) of *Synechococcus*-like A/B-lineage MAGs adding to existing literature on differentiation within this lineage. Using phylogenetic trees constructed from aligned PsaA and RbcL sequences, we found that six A/B-lineage MAGs were most closely related to A’-like ecotypes previously found in YNP and Hunter’s Hot Springs. These A’-like MAGs, which were recovered from the highest temperatures in our samples, have not been successfully cultured or fully sequenced to date, and recovery of multiple MAGs provides new potential to uncover adaptations to the thermal limits of photosynthesis. We found that within the samples tested, both temperature and total Fe controlled the range of A/B-lineage MAGs, while the pH range of A’-like MAGs was narrowed in comparison to the two other clades, suggesting that pH limits adaptations to high temperatures within these organisms. We uncovered genes that were enriched or depleted in each clade of A/B-lineage MAGs. Specifically, genes encoding both the light-independent protochlorophyllide reductase (DPOR) and nitrogenase were entirely absent from A’-like MAGs, suggesting that A’-like organisms must rely on fixed nitrogen from their environment, either from the overflowing water or from other diazotrophs in the community. Finally, we uncovered genes that diverge between clades of the A/B-lineage, particularly Cyt *c*_M_, which provides an avenue for future work into the regulation of photosynthesis under thermal stress and nutrient limited conditions in these organisms. Our work serves as an entry point for future hypothesis-based approaches to uncover physiological distinction between YNP hot spring cyanobacterial taxa.

## Materials and Methods

### Sample Collection, Sequencing, Metagenome Assembly, and Binning

Eleven sites throughout Yellowstone National Park were sampled with 16 total samples collected for metagenomic analysis. Sample temperatures ranged from 44°C to 73°C, pH ranged from 7.3 to 9.4. Generally, border between pigmented mats or filaments and the unpigmented chemotrophic zone was targeted for sequencing, with a variety of microbial structures sampled, thin and relatively thick mats to filaments. Samples for DNA extraction and water analysis were collected as previously described (Hamilton et al., 2019). Briefly, pH and temperature, were measured using a WTW pH 3310 meter (Xylem Analytics, Weilheim, Germany). A YSI 30 conductivity meter (YSI Inc, Yellow Springs, OH, USA) was used for conductivity measurements. A DR1900 portable spectrophotometer (Hach Company, Loveland, CO), was used to measure Fe^2+^, sulfide, and dissolved silica in overlaying water on-site. Cation concentration and trace element concentration were measured from 0.2-µm-filtered water as previously described (Hamilton & Havig, 2017, 2018; Havig et al., 2015).

Total DNA was extracted from triplicate ∼250-mg samples using a DNeasy PowerSoil kit (QIAGEN, Carlsbad, CA, USA). Triplicate DNA samples were pooled and sequenced on an Illumina HiSeq 2500 platform at the University of Minnesota Genomics Center (UMGC). Sickle (v. 1.33) was used to trim reads using PHRED score > 20 and minimum length > 50 as criteria. *De novo* assembly of short reads was performed on a per-sample basis using metaSPAdes (within SPAdes v. 3.11.0) for all samples except SM1 and SM2, which, due to high short read yield and computational limitations, were assembled using the Ray assembler (v. 2.3.1). Contigs were binned into MAGs using MetaBAT2 (v. 2.12.1).

### Metagenome Assembled Genome Quality Assessment, Refinement, and Taxonomic Abundance Estimation

The CheckM (v. 1.1.3) lineage workflow (Parks et al., 2015) was used to assess completeness, contamination, and strain heterogeneity of all MAGs collected via binning. For MAGs with greater than 10% contamination, manual coverage-based refinement was performed. Briefly, contigs with low coverage were removed, or in cases where two distinct clusters of contigs formed when GC and coverage were plotted, bins were split into separate MAGs by coverage. Refined bins were reassessed with CheckM.

Taxonomic classification was determined by pplacer (Matsen et al., 2010) and/or fastANI (Jain et al., 2018) within the Genome Taxonomy Database Toolkit (GTDB-Tk, v. 1.3.0) (Chaumeil et al., 2019), using the classify workflow and database version 89. The percent abundances of recovered MAGs was estimated using read mapping with BBMap. (v. 38.82) (Bushnell, 2015), adapted from a reads per kilobase of genome per gigabase of metagenome (RPKG) method described previously (Alcorta et al., 2020; Reji & Francis, 2020). Briefly the percentage of sample reads mapped to each MAG were normalized to genome size, then divided by the sum of normalized read abundances and multiplied by the ratio of mapped reads to total reads within a sample.

The formula used to calculate percent abundances was

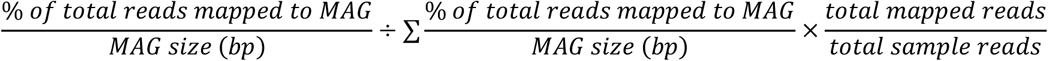

or briefly,

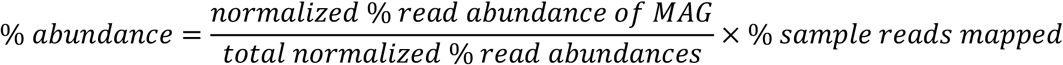

The results of this method are directly proportional to RPKG, except that they sum to 100. This was chosen because our conclusions were primarily drawn from intra-sample relative abundances. Furthermore, low overall RPKG values in some samples compared to others limited the ability to interpret intra-sample relative abundance. For comparison, an RPKG plot corresponding to Fig 2 can be found in Supplemental Material (Fig S3).

### Phylogenomic analysis

A set of 71 bacterial single copy marker genes present in each of 28 high quality (> 70% completeness, < 10% contamination) cyanobacterial MAGs and additional reference genomes was aligned and concatenated within anvi’o software (v. 7.1) (Eren et al., 2020). Open reading frames were identified within anvi’o using prodigal (Hyatt et al., 2010). From concatenated protein sequences, a phylogenomic tree was constructed using RAxML (Stamatakis, 2014) (100 bootstraps, protein gamma model automatically selected).

### Pangenome analysis and functional annotation

Pangenomes for MAGs predicted as A/B-like *Synechococcus, Gloeomargarita*, and Psuedanabaenaceae clades were constructed, visualized and analyzed using the anvi’o pangenomics workflow as previously described (Delmont & Eren, 2018), including closely related reference genomes in the analysis where possible. Gene clusters were predicted by first calculating open reading frame similarities using blastp (Altschul et al., 1990), then using the MCL algorithm (Dongen & Abreu-Goodger, 2012) to identify clusters within the blastp results. The inflation parameter for MCL was set to 10, to allow for high sensitivity in gene clustering across closely related genomes and decrease the occurrence of paralogs within clusters. Hierarchical clustering of gene clusters was performed using Euclidean distance and Ward linkages, and gene clusters were ordered in pangenome visualizations according to frequency. For each MAG and reference genome in a pangenome, gene functions were predicted using a BLAST search. NCBI COGs (2020 release), EGGNOGs, KEGG modules and KOfams were also predicted within anvi’o. MAGs and reference genomes in the *Synechococcus*-like pangenome were ordered based on maximum likelihood trees calculated with RAxML, using 100 bootstraps and the PROTGAMMA variable set to auto. MAGs were grouped into taxonomic clusters within anvi’o, which was then used to calculate enrichment of gene clusters with predicted functions. Additionally, KEGG module completeness was calculated for each MAG within anvi’o using the script anvi-estimate-metabolism.

### Functional homogeneity analysis

Gene cluster summaries were exported from the *Synechococcus*-like pangenome, which included aligned sequences, annotations, paralogs per genome, and function / geometric homogeneity scores. Gene clusters were then filtered in R by setting the max number of paralogs to 1, maximum functional homogeneity of 0.85, and minimum geometric homogeneity to be greater than functional homogeneity. Sequences from the resulting filtered gene clusters were extracted from the pangenome, aligned using Clustal Omega (Sievers & Higgins, 2014), and used to generate distance matrices and trees with RAxML (Stamatakis, 2014). Gene clusters that were diverged in the A’-clade from the other clades, were then selected by curation, using distance heatmaps as a guide.

## Acknowledgements

The authors would like to acknowledge that the research conducted for this work was done in Yellowstone National Park, which was created from land stolen from multiple Native American Nations, especially the Tukudeka (as well as other Shoshone-Bannock and Eastern Shoshone peoples) and Apsáalooke (Crow). These acts were done in part through the guise of Article 2 of the 1868 Fort Bridger Treaty and Article 2 of the 1868 Fort Laramie Treaty. The authors support efforts to give the lands encompassing YNP back to the native peoples who call it home. This research was supported by NASA Exobiology award number 80NSSC20K0614. T.L.H. conducts field research under permit YELL-2018-SCI-7020 issued to T.L.H. and Jeff Havig and reviewed annually by the Yellowstone Research Permitting Office. We are thankful to the Yellowstone Research Permitting Office, especially Annie Carlson and Erik Oberg, for provision of research permits and assistance in the process. We acknowledge the support of Minnesota Supercomputing Institute for providing crucial computing resources and data storage. We thank C. Grettenberger, A. Czaja, A. Gangidine, J. Miller, L. Penrose, L. Brengman, T. Djokic, A. Rutledge, L. Seyler, and J. Kuether for technical assistance in the field and A. Borowski for assistance processing samples in the lab.

## Data availability

Metagenome-assembled genomes have been deposited under NCBI BioProject PRJNA807728, and BioSamples SAMN26200794 to SAMN26200809. Full metagenome assemblies have been deposited to the Joint Genome Institute Integrated Microbial Genomes and Microbiomes database under accession numbers listed in Table 1.

---

**Figure S1.**
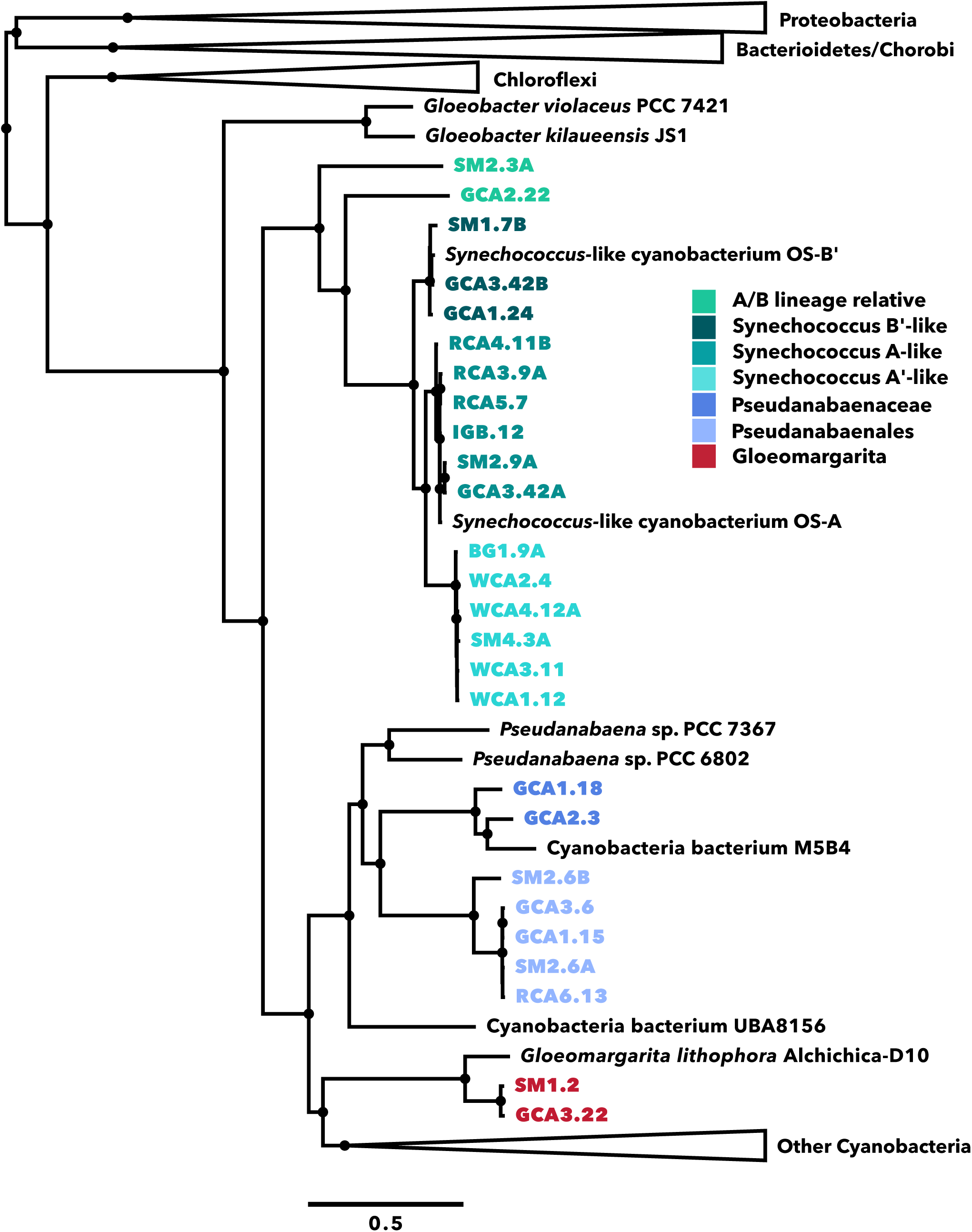
Phylogenomic tree of cyanobacterial MAGs. Distances were calculated from an aligned, concatenated set of 71 conserved marker genes from 26 high quality (> 70% completeness and < 10% contamination) cyanobacterial MAGs and a set of reference genomes. Node symbols denote bootstrap values above 80 from 100 bootstraps within RAxML.

**Fig S2.**
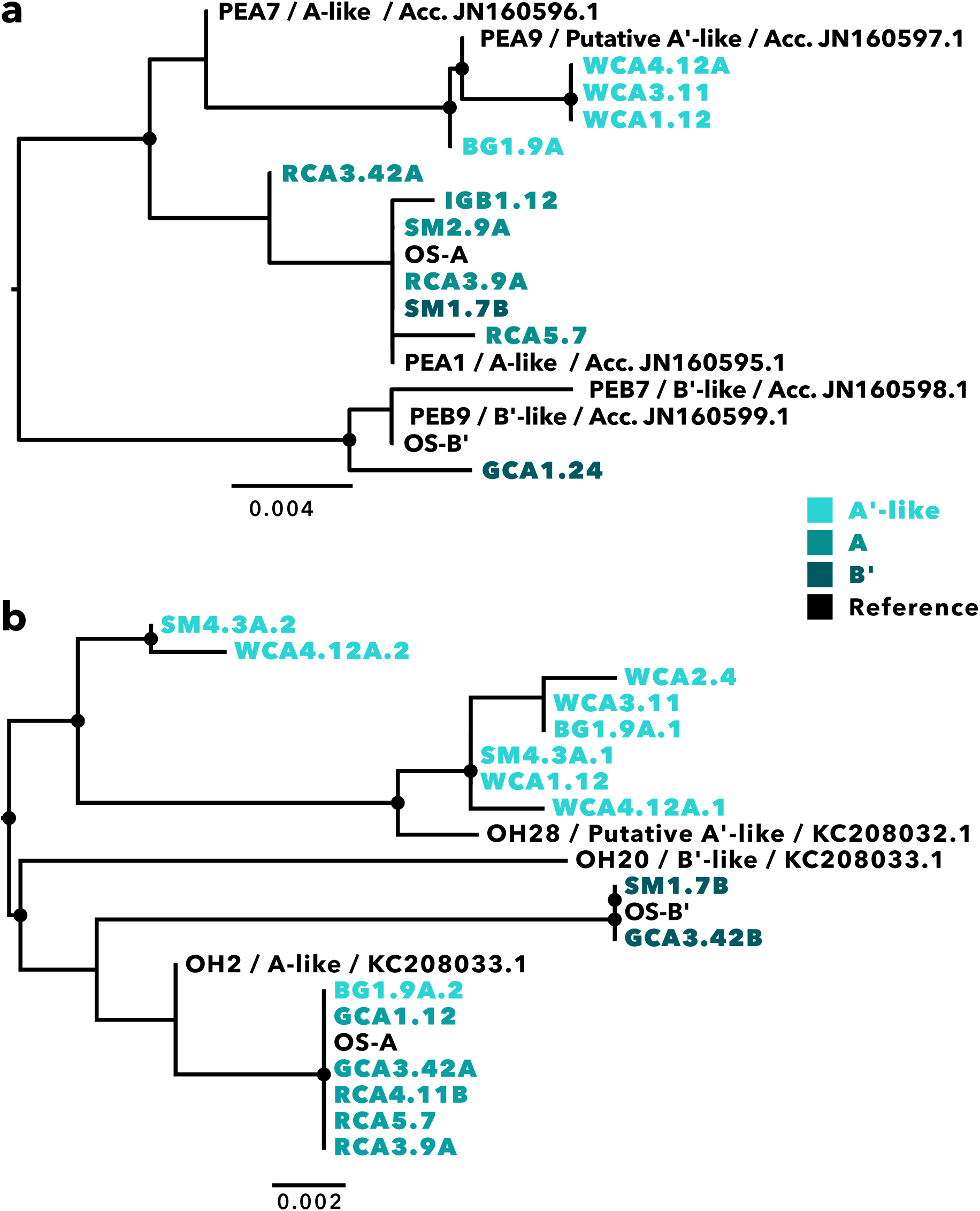
Support for taxonomic assignment of A’-like MAGs. **(a)** PsaA amino acid sequences extracted from 11 high-quality A/B-lineage MAGs compared with a subset of those published in Becraft et al., 2011. **(b)** 16 RbcL amino acid sequences extracted from 13 A/B-lineage MAGs compared to a subset published in Miller et al., 2013. Node symbols denote bootstrap values above 80 from 100 bootstraps within RAxML. Each sequence is named and colored according to MAG and taxonomic assignment. OS-A, OS-B’ and other published PsaA and RbcL sequences are in black.

**Table S1.** Proteins encoded by 71 single copy marker genes used for phylogenomic analysis

| Protein encoded |  |  |  |
| --- | --- | --- | --- |
| ADK | RecO_C | Ribosomal_L29 | Ribosomal_S6 |
| AICARFT_IMPCHas | Ribonuclease_P | Ribosomal_L3 | Ribosomal_S7 |
| ATP-synt | Ribosom_S12_S23 | Ribosomal_L32p | Ribosomal_S8 |
| ATP-synt_A | Ribosomal_L1 | Ribosomal_L35p | Ribosomal_S9 |
| Adenylsucc_synt | Ribosomal_L13 | Ribosomal_L4 | RsfS |
| Chorismate_synt | Ribosomal_L14 | Ribosomal_L5 | RuvX |
| EF_TS | Ribosomal_L16 | Ribosomal_L6 | SecE |
| Exonuc_VII_L | Ribosomal_L17 | Ribosomal_L9_C | SecG |
| GrpE | Ribosomal_L18p | Ribosomal_S10 | SecY |
| Ham1p_like | Ribosomal_L19 | Ribosomal_S11 | SmpB |
| IPPT | Ribosomal_L2 | Ribosomal_S13 | TsaE |
| OSCP | Ribosomal_L20 | Ribosomal_S15 | UPF0054 |
| PGK | Ribosomal_L21p | Ribosomal_S16 | YajC |
| Pept_tRNA_hydro | Ribosomal_L22 | Ribosomal_S17 | eIF-1a |
| RBFA | Ribosomal_L23 | Ribosomal_S19 | ribosomal_L24 |
| RNA_pol_L | Ribosomal_L27 | Ribosomal_S2 | tRNA-synt_1d |
| RNA_pol_Rpb6 | Ribosomal_L27A | Ribosomal_S20p | tRNA_m1G_MT |
| RRF | Ribosomal_L28 | Ribosomal_S3_C |  |

**Fig S3.**
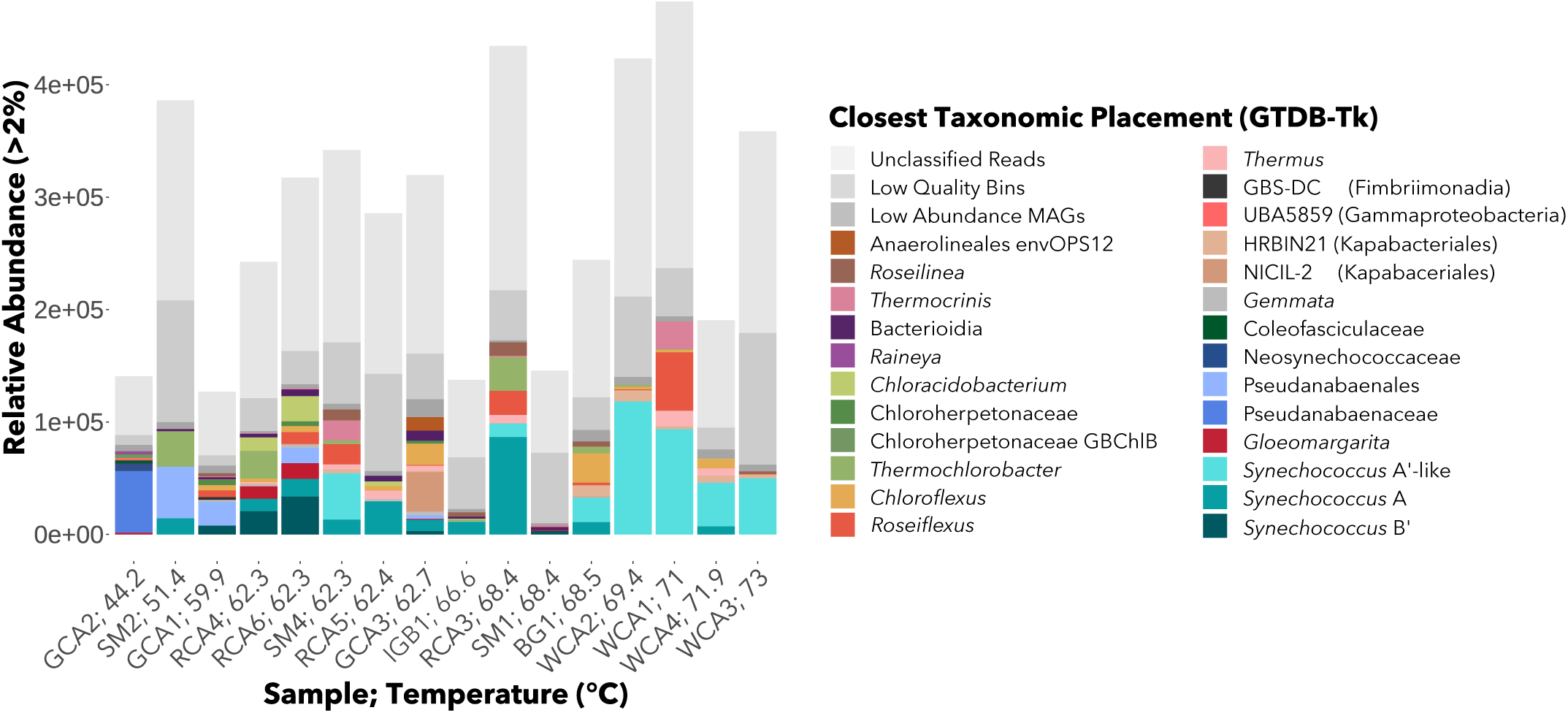
RPKG by temperature. RPKG of all MAGs with > 2% relative abundance in at least one sample. Unmapped reads, low quality bins, and low abundance MAGs are grouped per sample.

**Table S2.**
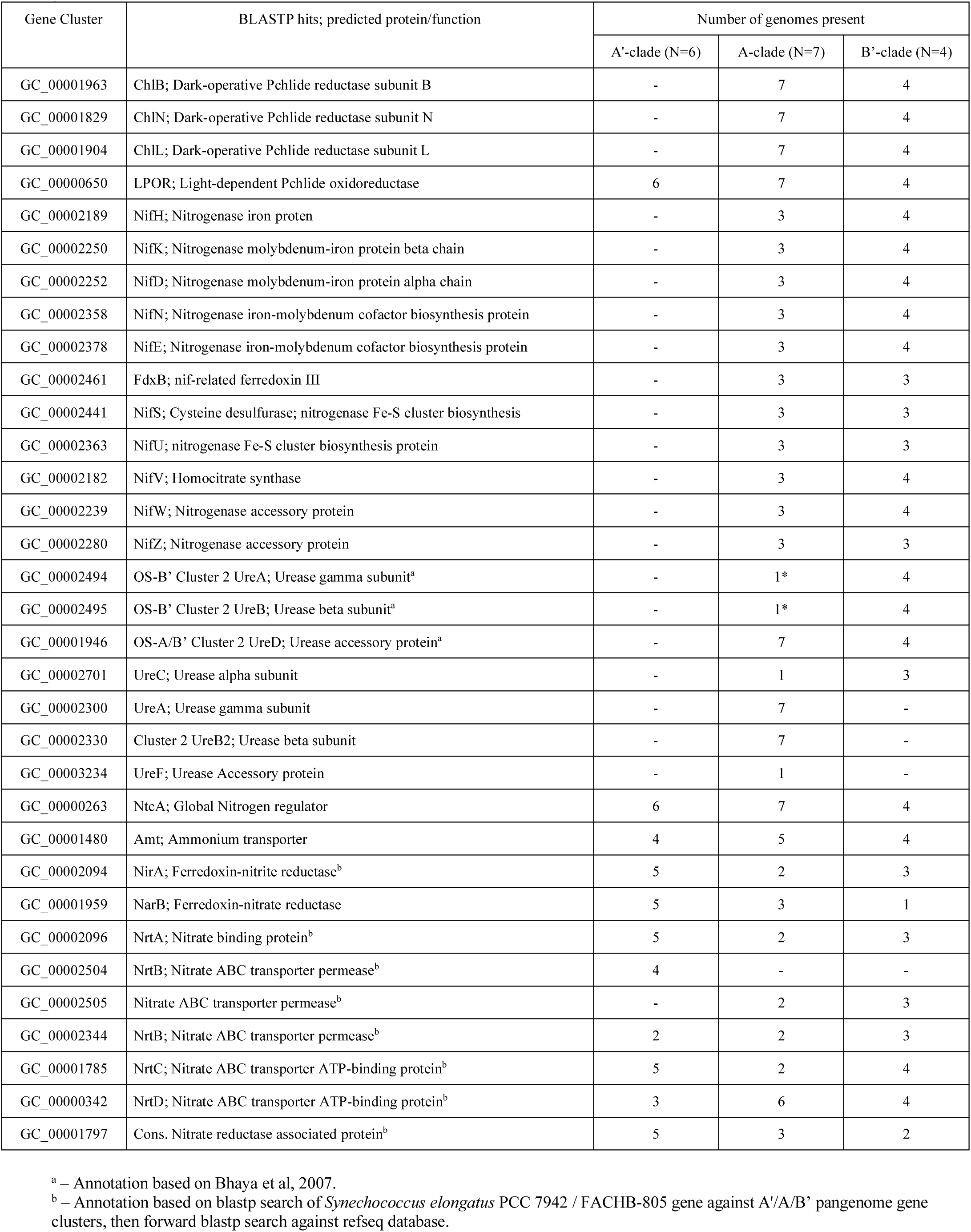
Nitrogen and protochlorophyllide reductase proteins encoded in A/B-lineage pangenome (15 MAGs, 2 reference genomes)

